# The apparent requirement for protein synthesis during G2 phase is due to checkpoint activation

**DOI:** 10.1101/863548

**Authors:** Sarah Lockhead, Alisa Moskaleva, Julia Kamenz, Yuxin Chen, Minjung Kang, Anay Reddy, Silvia Santos, James E. Ferrell

## Abstract

Protein synthesis inhibitors (e.g. cycloheximide) prevent cells from entering mitosis, suggesting that cell cycle progression requires protein synthesis until right before mitotic entry. However, cycloheximide is also known to activate p38 MAPK, which can delay mitotic entry through a G2/M checkpoint. Here we asked whether checkpoint activation or a requirement for protein synthesis is responsible for the cycloheximide effect. We found that p38 inhibitors prevent cycloheximide-treated cells from arresting in G2 phase, and that G2 duration is normal in about half of these cells. The Wee1/Myt1 inhibitor PD0166285 also prevents cycloheximide from blocking mitotic entry, raising the possibility that Wee1 and/or Myt1 mediate the cycloheximide-induced G2 arrest. Thus, the ultimate trigger for mitotic entry appears not to be the continued synthesis of mitotic cyclins or other proteins. However, M-phase progression was delayed in cycloheximide-plus-kinase-inhibitor-treated cells, emphasizing the different requirements of protein synthesis for timely entry and completion of mitosis.

**Impact statement:** Cycloheximide arrests cells in G2 phase due to activation of p38 MAPK, not inhibition of protein synthesis, arguing that protein synthesis in G2 phase is not required for mitotic entry.

## Introduction

Early studies on human cells in tissue culture as well as cells in the intestinal crypt of rats demonstrated that protein synthesis inhibitors like cycloheximide and puromycin prevent cells from entering mitosis, unless the cells were already in late G2 phase at the time of treatment (1, 2). The discovery of mitotic cyclins, activators of the cyclin-dependent kinases (Cdk), that accumulate prior to mitosis, provided a plausible explanation for these observations (3–5). Indeed, supplementing a cycloheximide-arrested *Xenopus* egg extract with exogenous cyclin B is sufficient to promote mitotic progression (6), as is supplementing an RNase-treated extract with cyclin B mRNA (7), and blocking the synthesis of cyclin B1 and B2 prevents mitotic entry (8). This argues that the synthesis of this particular protein is of singular importance for M-phase initiation.

In human cells, mitotic cyclins, mainly cyclins A2, B1, and B2, start to accumulate around the time of the G1/S transition as a result of the activation of cyclin transcription by E2F-family transcription factors (9) and stabilization of the cyclin proteins via APC/C^Cdh1^ inactivation (10). At the end of S phase, the ATR-mediated DNA replication checkpoint is turned off, and a FOXM1-mediated transcriptional circuit is activated (11). At about the same time, the pace of cyclin B1 accumulation (12–16), as well as the accumulation of other pro-mitotic regulators, including Plk1, Bora, and Aurora A, increases (12, 17, 18). These changes in transcription and protein abundances are thought to culminate in the activation of mitotic kinases, especially Cdk1, and the inactivation of the counteracting phosphatases PP1 and PP2A-B55 (19, 20). Cdk1 activity – judged by substrate phosphorylation – rises throughout G2 phase (12, 21) and sharply increases towards the end of G2 phase (12, 22). Cdk1-cyclin B1 then translocates from the cytoplasm to the nucleus just prior to nuclear envelope breakdown (16, 23–26).

The final increase in cyclin B1-Cdk1 activity, and decrease in PP2A-B55 activity, is thought to be due to the flipping of two bistable switches. Two feedback loops, a double-negative feedback loop involving the Cdk1-inhibitory kinases Wee1/Myt1 and a positive feedback loop involving the Cdk1-activating phosphatase Cdc25, keep Cdk1 activity low until cyclin B1 has reached a threshold concentration, beyond which the system switches from low to high Cdk1 activity, and high to low Wee1/Myt1 activity (Figure 1) (27–29). At the same time, a double negative feedback loop centered on PP2A-B55 flips and leads to an abrupt decrease of PP2A-B55 activity (30–34).

**Figure 1.**
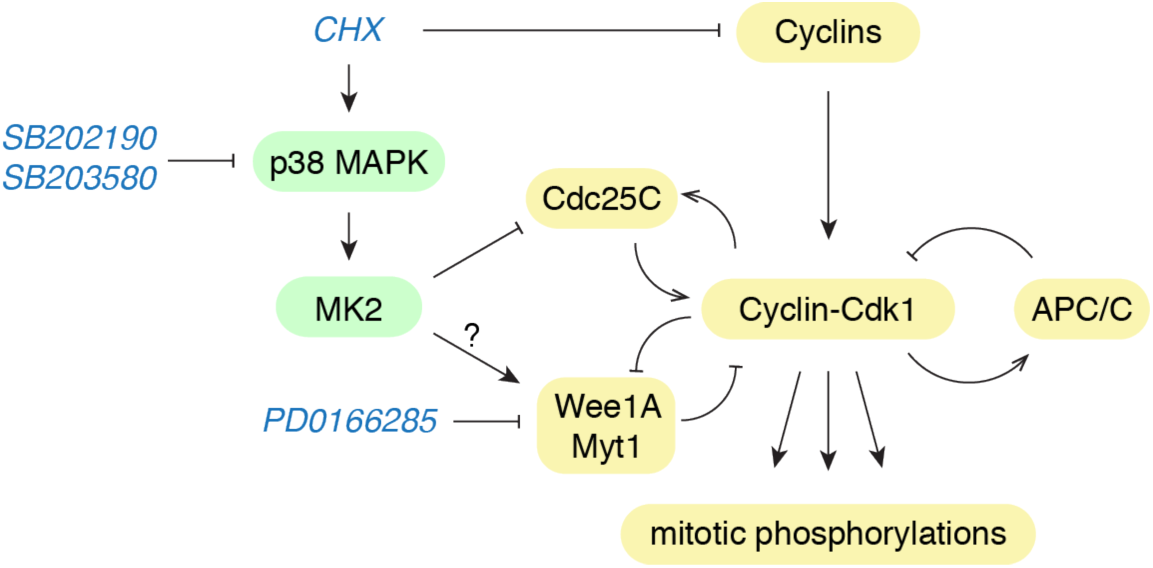
The mitotic entry network. Schematic of the regulation of Cdk1 activity at the G2/M transition by cyclins and multiple feedback loops. The protein synthesis inhibitor cycloheximide (CHX) can block cyclin accumulation, and activate p38 MAPK causing a delay in G2/M progression by inhibiting Cdc25 and/or potentially activating Wee1/Myt1 (36).

The cyclin B1 threshold concentration is determined by the amounts of Cdc25 and Wee1/Myt1 activity present (35). In somatic cells, several signaling pathways impinge upon Cdc25 and/or Wee1 to delay the G2-to-M transition in the face of stresses (36). These include the ATM/ATR kinases, which activate Chk1 and Chk2, which in turn can inactivate Cdc25 and activate Wee1 by phosphorylating 14-3-3 binding sites in the two Cdk1 regulators. These pathways play a role in delaying mitosis in the presence of DNA damage, and may also help prevent premature mitosis in cells undergoing normal DNA replication (36–38). In addition, a protein kinase cascade that includes the MKK3 and MKK6 MAP kinase kinases, p38 MAPK, and the downstream kinase MAPKAP kinase 2 (MK2), has been implicated in the negative regulation of Cdc25, by phosphorylating the same 14-3-3 binding site (39). Interestingly, p38 activation has been observed in response to protein synthesis stresses, including cycloheximide (40); indeed cycloheximide is often used as a positive control for maximal activation of p38. These findings raise the question of whether the cycloheximide-dependent G2 delay is indeed caused by blocking the synthesis of proteins required for mitotic entry, or rather activation of the p38-dependent G2/M checkpoint.

Here we used live-cell markers of cell cycle progression combined with small molecule inhibitors to dissect the contribution of protein synthesis to G2 and mitotic progression. We show that inhibition of Wee1/Myt1 shortens the duration of G2 phase in a dose-dependent manner, and allows cells to progress into mitosis in the presence of cycloheximide. Moreover, p38 inhibition overcomes a cycloheximide-induced G2 arrest, arguing that p38-mediated checkpoint activation causes the arrest and not insufficient protein synthesis. However, although G2 protein synthesis was not required for mitotic entry, it was required for normal mitotic progression. These findings suggest that the burst of cyclin synthesis that normally occurs during G2 phase serves as a “just-in-time” preparation for mitotic progression, but does not trigger mitotic entry.

## Results

We chose MCF10A cells, a spontaneously immortalized human mammary epithelial cell line, for these studies, because they are euploid, non-tumorigenic, and have been studied extensively (41, 42). To determine when S phase ends and G2 phase begins, we stably expressed an eYFP-PCNA fusion protein, a live-cell marker of DNA replication (43, 44). eYFP-PCNA forms bright foci within the nucleus during S phase, which become brighter and less numerous as S phase progresses (Figure 2A,B). We verified that eYFP-PCNA foci co-localized with BrdU and EdU-staining sites of active DNA, as previously reported (44). At the end of S phase, eYFP-PCNA foci dissolve and fluorescence becomes diffuse, marking the S/G2 transition (Figure 2B). Upon nuclear envelope breakdown (NEB), nuclear eYFP-PCNA disperses throughout the cell; this can be taken as a marker for the G2/M transition (or, more precisely, of the prophase/prometaphase transition; Figure 2A). Thus eYFP-PCNA proved to be well suited to measure G2 duration, from the time of G2 onset (the disappearance of foci) to the time of G2 termination (taken as the time when eYFP-PCNA exited the nucleus due to NEB). Typical mean G2 durations were about 4 h with a variance of 25%, within the range of previously reported durations for G2 phase in a variety of cell lines (12, 43, 45–48).

**Figure 2.**
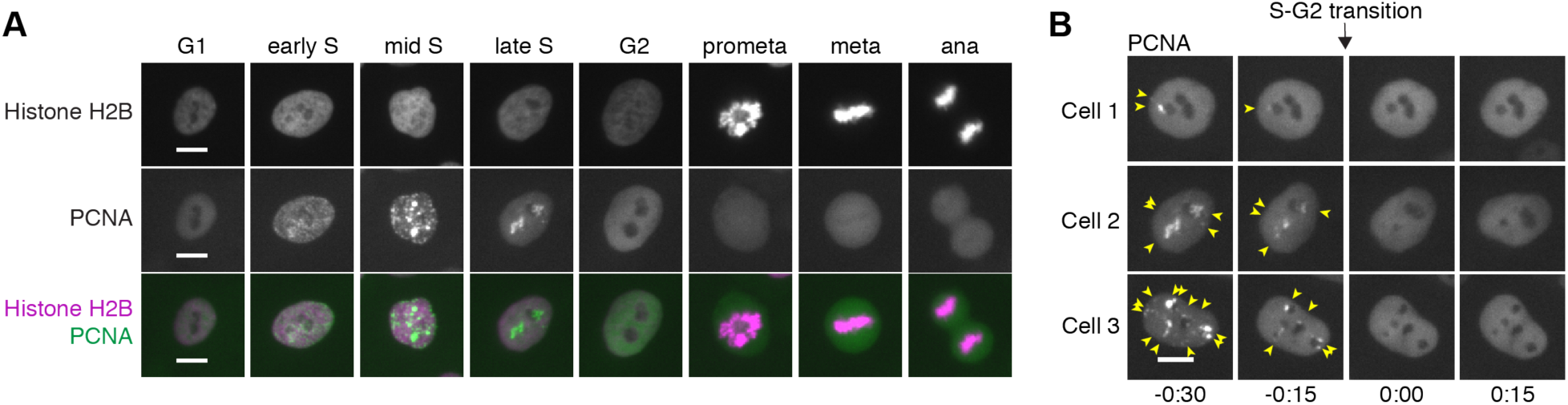
Measuring the duration of cell cycle phases using fluorescently labelled PCNA and histone H2B in MCF10A cells. (**A**) eYFP-PCNA can be used to determine the onset of S phase, the completion of S phase, and the onset of mitosis (nuclear envelope breakdown); histone H2B-mTurquoise (used here) or histone H2B-mCherry can be used to determine anaphase onset (**B**) Three examples of cells showing the disappearance of eYFP-PCNA foci (arrows) at the end of S phase. Time (in the format h:min) was aligned to the time of entry into G2 phase. Scale bars: 10 *µ*m.

### Live-cell imaging confirms that cycloheximide blocks entry into mitosis

Early studies on fixed cells showed that the protein synthesis inhibitors puromycin and cycloheximide cause cells to arrest in G2 phase (1, 2). We confirmed this finding by live-cell microscopy using the PCNA probe to demarcate G2 phase. We followed asynchronously growing cells in cell culture for 4-6 h, then added cycloheximide (10 *µ*g/ml) and continued to follow the cells for another 6-10 h (Figure 3A). This allowed us to identify cells which had exited S phase during the initial imaging period, determine accurately how much time these cells had spent in G2 phase prior to drug addition, and finally determine the fate of these cells in response to the drug treatment. Because many of our subsequent experiments required the addition of DMSO-solubilized drugs to a final DMSO concentration of 0.1%, we performed all experiments in the presence of this concentration of DMSO.

**Figure 3.**
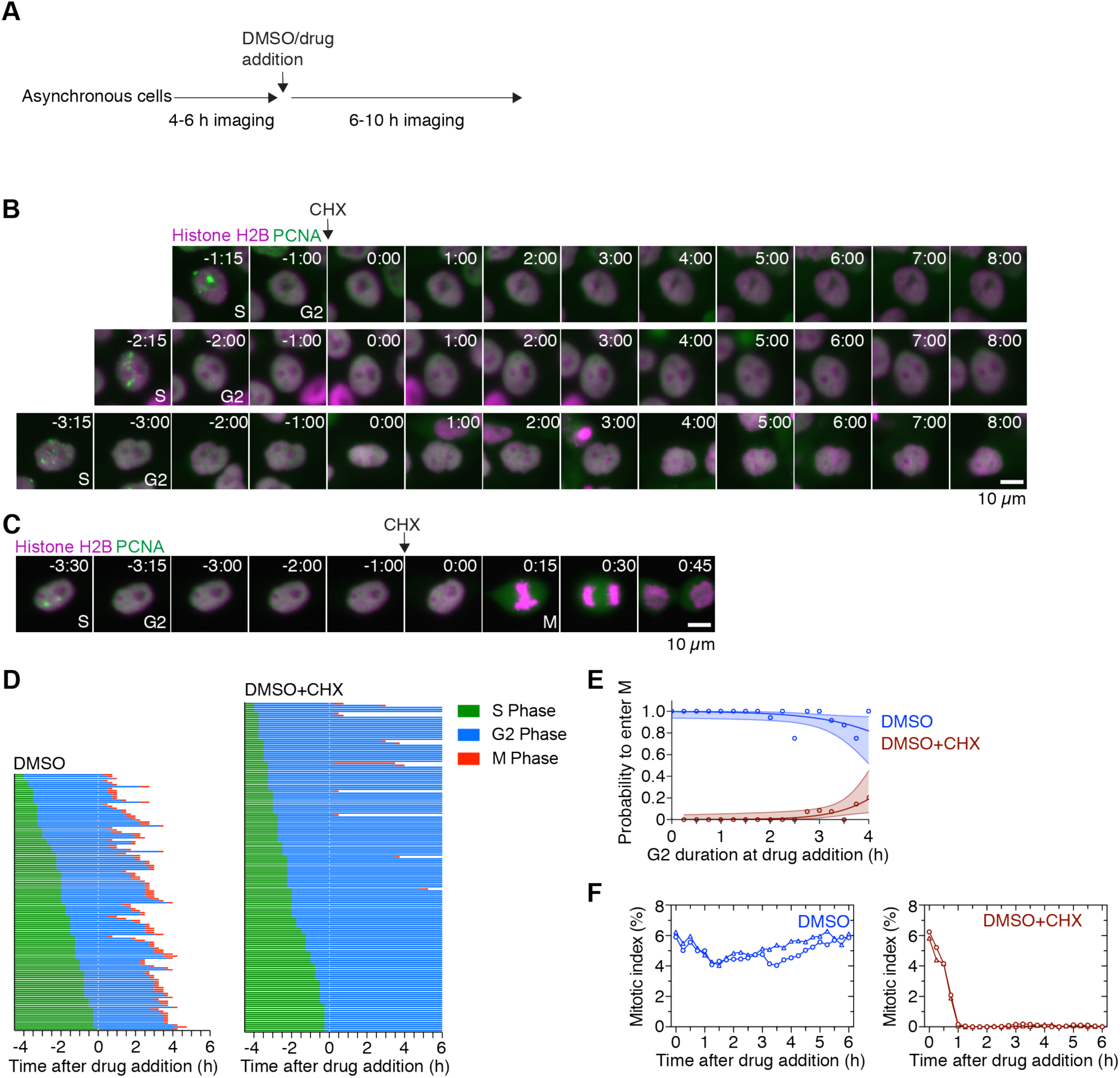
Cycloheximide blocks entry into mitosis. (**A**) Schematic of the experimental setup: asynchronously grown cells were imaged for 4-6 h in order to determine the time when cells exited S phase. After this period DMSO or small molecule inhibitors were added, and cells were followed for another 6-10 h to determine whether and when the cells entered mitosis. (**B-C**) Montages of MCF10A cells expressing H2B-mCherry and eYFP-PCNA followed over the time course of the experiment described in (A). Time (in the format h:min) was aligned to the point of DMSO or cycloheximide addition. Three cells are shown that had spent different amounts of time in G2 phase at the time of cycloheximide addition, and subsequently either arrested in G2 phase (B) or entered mitosis (C). (**D**) Cell cycle progression in MCF10A cells expressing H2B-Turquoise and eYFP-PCNA and treated with DMSO (left, n=130) or CHX (right, n=165). (**E**) Logistic regression analysis. This estimates the probability of a cell entering mitosis as a function of how long the cell had been in G2 phase at the time of drug addition, for the experiment shown in (D). Circles indicate the fraction of cells that ultimately entered mitosis for given times of drug addition. The solid lines show the logistic fit for the data and the lightly colored areas indicate the 95% confidence intervals. (**F**) Mitotic indices for MCF10A cells expressing H2B-Turquoise and eYFP-PCNA cells treated with DMSO or CHX. Shown are two independent experiments (circles and triangles, respectively). At least 3605 cells were counted for each timepoint.

Cells treated with DMSO alone progressed into mitosis (130 out of 130 cells), but cycloheximide addition arrested the large majority of cells (153 cells out of 165) in G2 phase (Figures 3B-D). Cycloheximide-treated cells were more likely to progress into mitosis if the drug was added late in G2 phase. Of the 12 cycloheximide-treated cells that did enter mitosis, 10 had spent more than 3 h in G2 phase (>75% of the duration of a normal G2 phase) at the time of cycloheximide addition (Figure 3C,D). Based on logistic regression analysis, the probability that a cycloheximide-treated cell will enter mitosis if the cycloheximide is added 2 h after the start of G2 phase is 1% (with a 95% confidence interval (CI) of 0 to 7%); if added 3 h after the start of G2 phase, it rises to 4% (95% CI 1 to 11%); and if it is added 4 h after the start of G2 phase, the duration of a typical normal G2 phase, the probability is 19 (95% CI 6 to 45%) (Figure 3E). The fraction of mitotic cells in the cell population (mitotic index) remained approximately constant throughout the experiment for the DMSO-treated population, but decreased to near-zero within 60 min after cycloheximide treatment (Figure 3F). Together, these findings confirm that cycloheximide-treated G2 cells do arrest, as previously noted (1, 2), and imply that cells remain sensitive to cycloheximide treatment until late in G2 phase.

### Wee1/Myt1 inhibition shortens G2 phase and restores mitotic entry in cycloheximide-treated G2 phase cells

The Wee1/Myt1 kinases are key regulators of the G2/M transition that, when active, restrain Cdk1-cyclin B activity. FRET studies have indicated that Cdk1 activation begins just prior to the nuclear translocation of Cdk1-cyclin B1—thus very late in G2 phase—which suggests that Wee1 and Myt1 may be on during almost all of G2 phase (22, 49). However, other studies have suggested that some Cdk1 activation can be detected early in G2 phase (12), which could mean that Wee1/Myt1 is switched off earlier. These possibilities can be distinguished by determining how much the duration of G2 phase can be shortened by Wee1/Myt1 inhibition. In the former case, the minimal duration of G2 phase would be near zero; in the latter case it would be longer, with the minimal duration of G2 phase corresponding to how long the interval normally is between the inactivation of Wee1/Myt1 and the onset of M phase.

To inhibit Wee1/Myt1 activity we used the small molecule inhibitor PD0166285 (50). Treating an asynchronously growing cell culture with different concentrations of PD0166285 reduced the Wee1-mediated phosphorylation of Cdk1 at Tyr 15 in a dose-dependent manner (Figure 4A).

**Figure 4.**
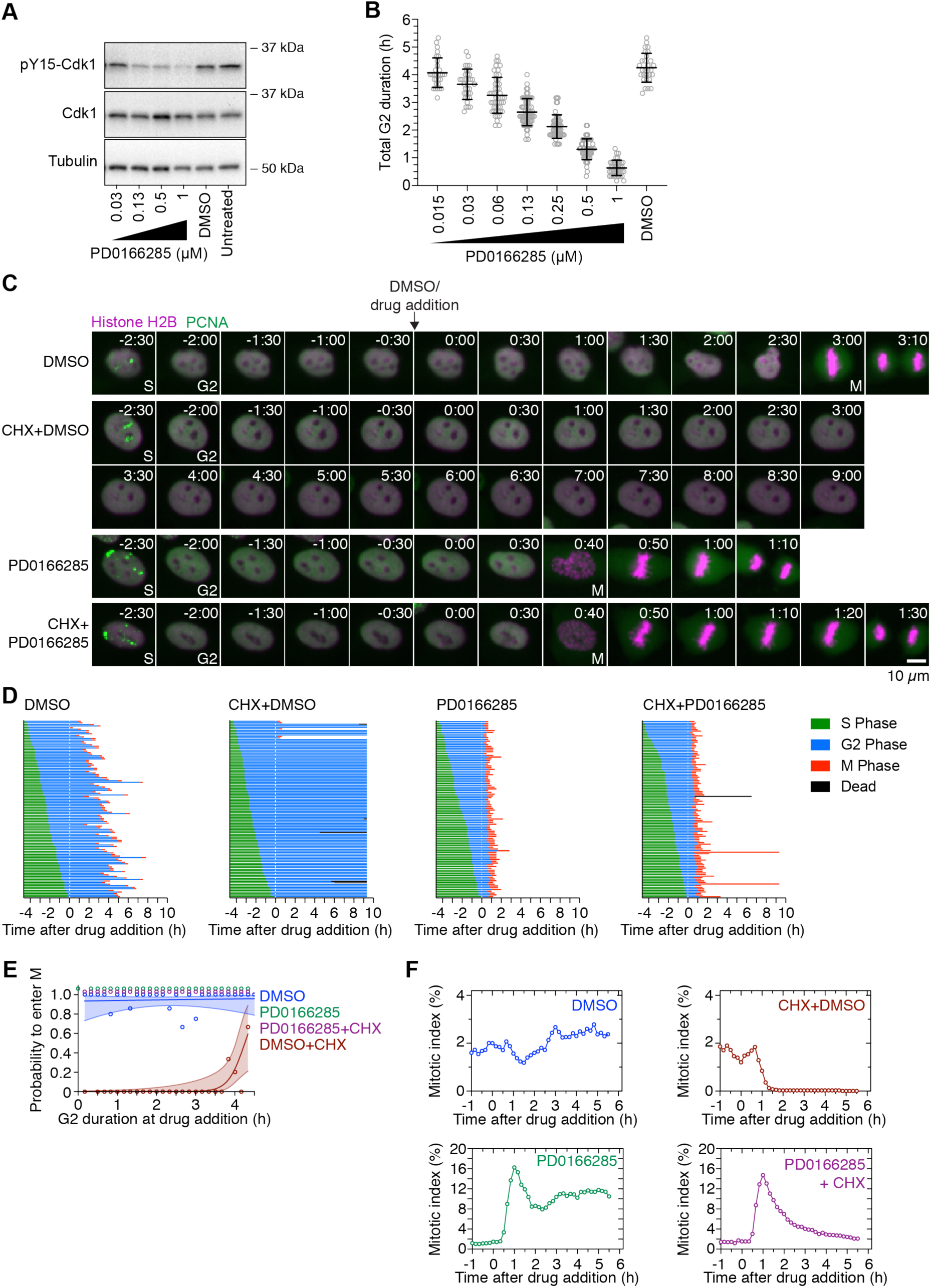
Wee1/Myt1 inhibition shortens G2 phase and restores mitotic entry in cycloheximide-treated G2-phase cells. (**A**) Asynchronously growing MCF10A cells were treated with DMSO or different concentrations of the Wee1/Myt1-inhibitor PD0166825 for 30 min and the phosphorylation state of tyrosine 15 of Cdk1 was analyzed by immunoblotting as a measure of Wee1/Myt1 activity. Cdk1 and *α*-tubulin were used as loading controls. (**B**) The length of G2 phase was measured by live cell fluorescence microscopy of cells expressing H2B-mCherry and eYFP-PCNA in the presence of DMSO or different concentrations of PD0166285. Only cells that had not entered G2 phase at the time of treatment and showed a distinct G2 phase were included in this analysis (n > 28 cells for all conditions). (**C**) Montages of MCF10A cells expressing H2B-mCherry and eYFP-PCNA followed over the time course of the experiment described in Figure 3A. Time (in the format h:min) was aligned to the point of DMSO/drug addition (10 *µ*g/mL cycloheximide (CHX) and/or 1 *µ*M PD0166285). (**D**) Cell cycle progression for MCF10A cells expressing H2B-mCherry and eYFP-PCNA and treated with DMSO (n=113), cycloheximide (n=116), PD0166285 (n=100) or cycloheximide plus PD0166285 (n=111). The majority of cells treated with cycloheximide arrested in G2 phase, while cells treated with 1 *µ*M PD0166825 alone or cycloheximide plus 1 *µ*M PD0166825 progressed into mitosis shortly after treatment with the drug. (**E**) Logistic regression analysis. Probability of a cell entering mitosis as a function of how long the cell had been in G2 phase at the time of drug addition for the experiment shown in (D). Circles indicate the fraction of cells that ultimately entered mitosis for given times of drug addition. The solid lines show the logistic fits for the data and the lightly colored areas indicate the 95% confidence intervals. Note that the green (PD0166285) and purple (PD0166285+CHX) data points have been shifted upward to make them visible. (**F**) Mitotic indices for MCF10A cells expressing H2B-mCherry and eYFP-PCNA treated with DMSO, CHX, PD0166285 or CHX plus PD0166285. At least 3226 cells were counted for each timepoint.

We next investigated the impact of Wee1/Myt1 inhibition on the duration of G2 phase by live-cell imaging. Cells were treated with different concentrations of PD0166285, and 28-60 cells that entered and completed G2 phase over 6 h of imaging were tracked and their G2 durations were determined (Figure 4B). PD0166285 shortened G2 phase in a graded fashion. The highest concentration of PD0166285 (1 *µ*M) resulted in a G2 duration of 38 ± 17 min compared to 255 ± 31 min in the DMSO-treated control (mean ± S.D., Figure 4B). This suggests that the Wee1/Myt1 switch is normally thrown very late in G2 phase. This also suggests that there is sufficient cyclin (and any other proteins essential for M phase entry) present even early in G2 phase to allow rapid mitotic entry, provided that Wee1/Myt1 activity is low. Moreover, the basal level of Wee1/Myt1 activity determines the length of G2 phase. These results are consistent with and extend the findings of previous studies on the effects of Wee1/Myt1 inhibition (45, 50, 51).

Considering the central role of Wee1/Myt1 in controlling G2/M, we asked whether Wee1/Myt1 inhibition was able to overcome the cycloheximide-induced G2 arrest. We used the same experimental set up as described in Figure 3A, but after the initial imaging period added DMSO, cycloheximide, 1 *µ*M PD0166285, or cycloheximide plus 1 *µ*M PD0166285 to the cells. Again (cf. Figure 3D) cells treated with DMSO progressed into mitosis with normal G2 duration whereas cycloheximide prevented most cells from entering mitosis (Figure 4C-E and Figure 4–figure supplement 1A). All but one (99/100) of the PD0166285-treated cells entered mitosis within one hour of drug addition (Figure 4C,D, and Figure 4–figure supplement 1A). Cells that were treated with PD0166285 late in G2 phase tended to enter mitosis more quickly than those treated early in G2 phase (Figure 4–figure supplement 1B). Consistent with these findings, the fraction of cells in mitosis spiked about 10-fold within the first hour of PD0166285 treatment and remained elevated for the rest of the experiment (Figure 4F).

Strikingly, cycloheximide did not block mitotic entry in the presence of PD0166285 (Figure 4C,D, and Figure 4–figure supplement 1A), and cells progressed into mitosis with similar dynamics as cells treated with PD0166285 alone (Figure 4–figure supplement 1B). In contrast to cycloheximide treatment alone, the probability of a cell entering mitosis was ∼100%, and was independent of the time the cell had spent in G2 phase at the time of drug addition (Figure 4E). This indicates that PD0166285 can override the cycloheximide-induced G2 arrest. The override was also observed when cells were treated with cycloheximide for 2 h prior to the addition of PD0166285 (Figure 4–figure supplement 1C). As we had observed for cells treated with the Wee1/Myt1 inhibitor alone, the mitotic index for cells treated with PD0166285 plus cycloheximide showed a pronounced spike within the first hour of treatment; however, the spike then slowly decayed over time (Figure 4F). The decay is consistent with the hypothesis that even though PD0166285 abrogates the need for protein synthesis in G2 phase cells, cells earlier in the cell cycle still do need protein synthesis to ultimately carry out mitosis.

### Cycloheximide treatment in S phase blocks cell cycle progression even in the absence of Wee1/Myt1 activity

Previous studies of Wee1 inhibitors have shown that they can drive chemotherapy-treated and p53-mutant cell lines into mitosis without completing DNA replication (52). To further explore the connection between protein synthesis and M-phase entry, we analyzed the cell cycle progression of cells treated with cycloheximide, PD0166825, or PD0166285 plus cycloheximide while still in S phase. After being treated with DMSO in S phase, 100% of cells (85/85) entered G2 phase and 84% (71/85) of these cells progressed into mitosis during the 10 h imaging period (Figure 5A). In contrast, only 33% of cells (32/98) treated with cycloheximide during S phase progressed into G2 phase, whereas all other cells continued to display PCNA foci (albeit dimmer foci than those seen in control cells), suggesting that S phase was never completed (Figure 5A,B). All cells (99/99) treated with 1 *µ*M PD0166285 during S phase entered mitosis; remarkably, 38 of them progressed into mitosis in the presence of PCNA foci and without displaying a detectable G2 phase (Figure 5A,C), suggesting that at some stage of S phase there is enough pro-mitotic activity to drive cells into mitosis if Wee1/Myt1 are inhibited. However, most of the cells that had directly progressed into mitosis failed to undergo proper chromosome segregation and cytokinesis (28/38 failures compared to 3/61 cells which displayed a G2 phase, Figure 5A,C) and even those cells that carried out some duration of G2 phase prior to mitotic entry required more time to progress through mitosis (Figure 5D).

**Figure 5.**
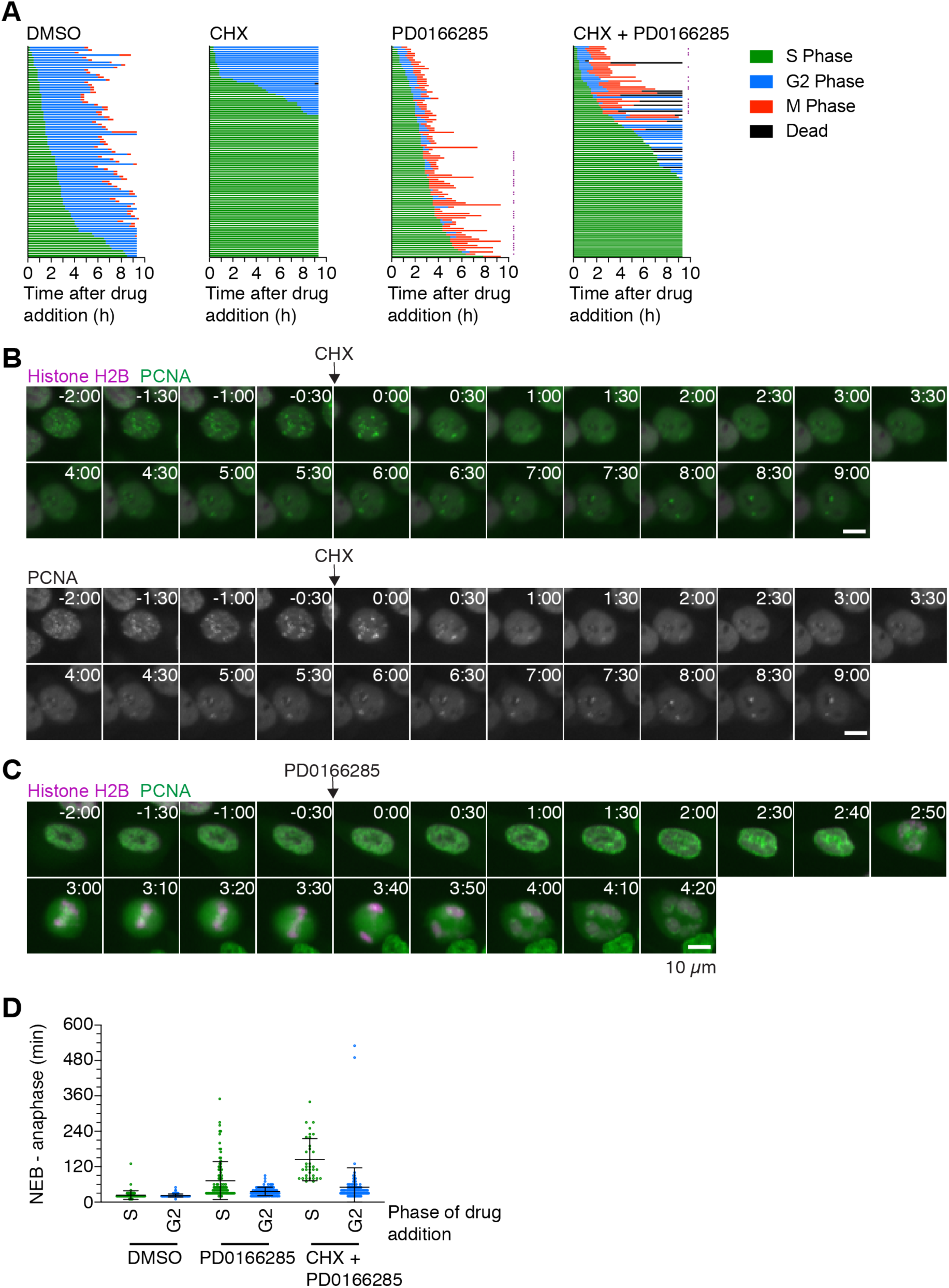
Cycloheximide treatment in S phase blocks cell cycle progression even in the absence of Wee1/Myt1 activity. (**A**) Cell cycle progression for MCF10A cells expressing H2B-mCherry and eYFP-PCNA treated during S phase with either DMSO (n=85), cycloheximide (n=98), 1 *µ*M PD0166285 (n=99), or cycloheximide plus 1 *µ*M PD0166285 (n=105). Cells marked with a purple square showed abnormal mitotic progression, often lacking proper metaphase and cytokinesis. (**B**) Montage of an MCF10A cell expressing H2B-mCherry and eYFP-PCNA treated with cycloheximide during S phase. In this cell (and most cells), the PCNA foci became weaker after cycloheximide treatment, yet they never completely disappeared, suggesting that these cells remained in S phase. Time (in the format h:min) was aligned to the point of cycloheximide addition. (**C**) Montage of an MCF10A cell expressing H2B-mCherry and eYFP-PCNA treated with 1 *µ*M PD0166285 during S phase that progressed into mitosis in the presence of PCNA foci (suggesting that the cell never completed S phase). Note the abnormal mitotic progression without a proper metaphase and cytokinesis. (**D**) Mitotic duration, measured as the time from nuclear envelope breakdown (NEB) to anaphase, for cells treated either in G2 phase or S phase with DMSO, PD0166285, or cycloheximide plus PD0166285. Whereas treatment with PD0166285 or treatment with PD0166285 plus cycloheximide in G2 phase only slightly extended mitosis (see also Figure 7), treatment with these drugs in S phase dramatically increased the duration of mitosis.

Whereas Wee1/Myt1 inhibition had been able to overcome a cycloheximide arrest when cells had already entered G2 at the time of drug addition, only 32% of cells (34/105) treated with cycloheximide plus PD0166285 in S phase entered mitosis, 9 of them directly from S phase. 33% of the cells (38/105) never completed S phase and 19% (20/105) entered G2 phase but not mitosis. Cells that managed to enter mitosis frequently exhibited extended and qualitatively abnormal mitotic progression (Figure 5A,D). These findings underscore the hypothesis that some S-phase protein synthesis is required for mitotic entry even in the absence of the Cdk1-inhibiting activity of Wee1/Myt1.

### Wee1/Myt1 counteract pro-mitotic activities that accumulate during G2 phase

To further investigate the relationship between Wee1/Myt1 activity, G2 duration and a cell’s ability to enter mitosis, we followed untreated cells for 6 h, treated cells with cycloheximide for 2 h, then added different Wee1/Myt1 inhibitor concentrations and assessed the cell’s fate (Figure 4–figure supplement 1C,D). At the lowest concentration of PD0166285 (0.125 *µ*M), only a fraction of cells (31%, 32/104) progressed into mitosis and the probability for a cell to enter mitosis increased as the time the cell had spent in G2 phase prior to cycloheximide addition increased (Figure 4–figure supplement 1D). With increasing PD0166285 concentrations, more cells were able to enter mitosis even if they had spent less time in G2 phase prior to drug addition (Figure 4–figure supplement 1C,D). These results are consistent with the idea that pro-mitotic activities accumulate throughout G2 phase and are opposed by Wee1/Myt1 activity during this time. The later in G2 phase, the more pro-mitotic activities have accumulated and the less completely Wee1/Myt1 needs to be inhibited in order to flip the mitotic switch.

### p38 inhibition allows cells to enter mitosis in the presence of cycloheximide

The data presented so far are consistent with the hypothesis that in G2 phase cyclin synthesis triggers mitotic entry: cycloheximide blocks mitotic entry, and the effects of PD0166285 suggest that some pro-mitotic activity, possibly cyclin, gradually accumulates throughout G2 phase. However, it also remains possible that the ability of cycloheximide to block mitotic entry is due to its activation of p38 MAPK and MK2, rather than to any effect on cyclin accumulation (36, 40).

To address this issue directly, we treated cells with cycloheximide plus one of two p38 MAPK inhibitors (SB202190 and SB203580) that act as high affinity inhibitors of p38α and p38β (MAPK14 and MAPK11) and as lower affinity inhibitors of other protein kinases (53). We verified that, as previously reported, cycloheximide stimulated the phosphorylation of p38 and its downstream target Hsp27, and that the inhibitors decreased cycloheximide-induced Hsp27 phosphorylation in a dose-dependent fashion (Figure 6A).

**Figure 6.**
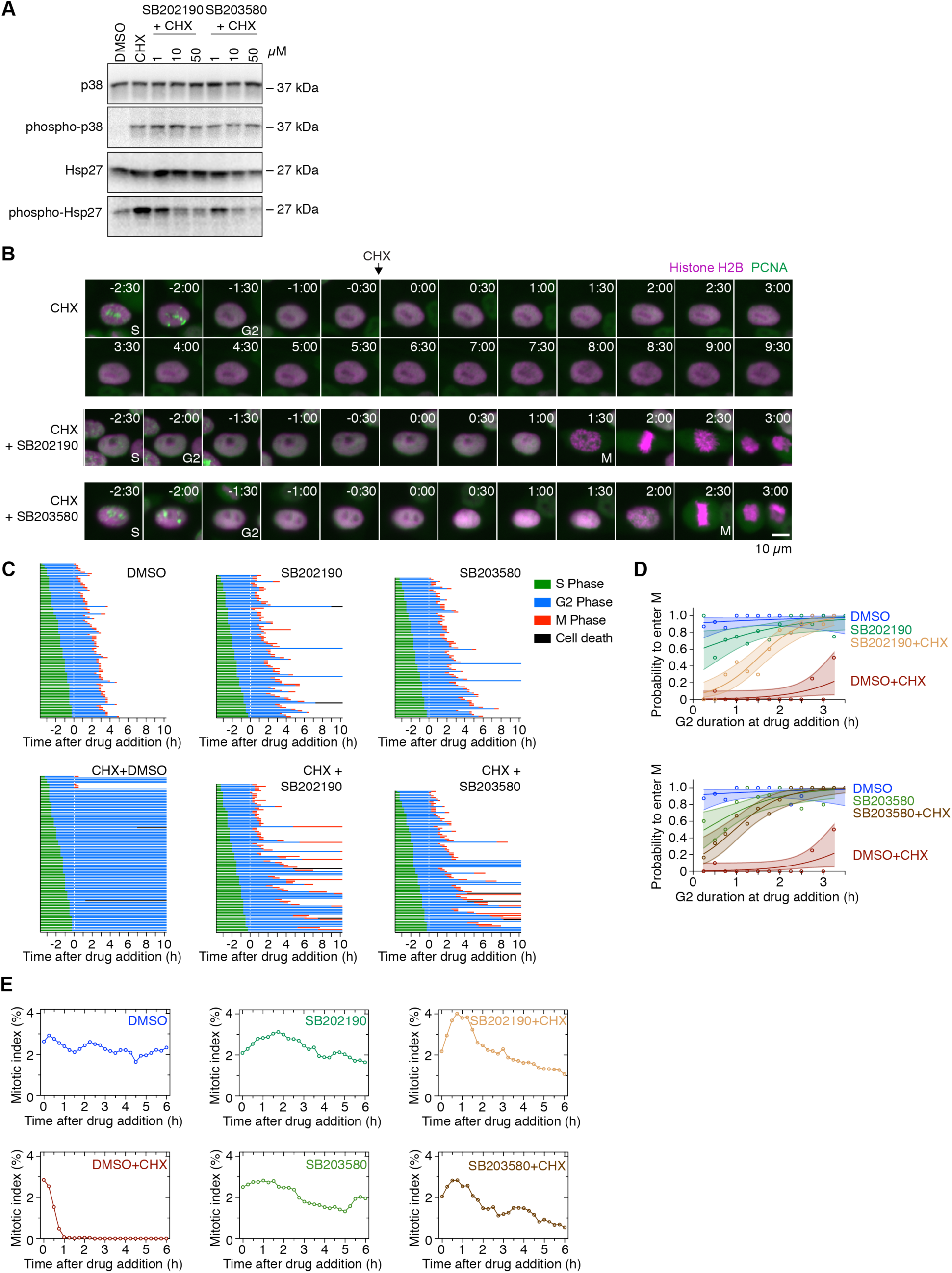
p38 MAPK inhibition allows cells to enter mitosis in the presence of cycloheximide. (**A**) Asynchronously growing cells were treated for 6 h with DMSO, cycloheximide or cycloheximide plus either of the p38 MAPK inhibitors SB202190 or SB203580. The phosphorylation state of p38 as well as the phosphorylation state of the p38 substrate Hsp27 was analyzed by immunoblotting to assess the activation state of p38. p38 and Hsp27 were used as loading controls. Whereas cycloheximide induced the phosphorylation of both p38 and Hsp27, CHX plus either SB202190 or SB203580 reduced the p38-mediated phosphorylation of Hsp27, but not the phosphorylation of p38 itself. (**B**) Montages of MCF10A cells expressing H2B-mCherry and eYFP-PCNA followed over the time course of the experiment described in Figure 3A treated with cycloheximide or cycloheximide plus either SB202190 or SB203580. Time (in the format h:min) was aligned to the point of drug addition. (**C**) Cell cycle progression for MCF10A cells expressing H2B-mCherry and eYFP-PCNA treated with DMSO (n=100), CHX (n=100), SB202190 (n=92), SB203580 (n=90), CHX+SB202190 (n=95) or CHX+SB203580 (n=91). The majority of cells treated with cycloheximide arrested in G2 phase, whereas cells treated with CHX plus SB202190 or SB203580 (50 *µ*M) progressed into mitosis in most cases. (**D**) Logistic regression analysis. Probability of a cell to enter mitosis as a function of how long the cell has already been in G2 phase at the time of drug addition for the experiment shown in (C). Circles indicate the fraction of cells that ultimately entered mitosis for given times of drug addition. The solid lines show the logistic fit for the data and the lightly colored areas indicate the 95% confidence intervals. (**E**) Mitotic indices for MCF10A cells expressing H2B-mCherry and eYFP-PCNA cells treated with DMSO, CHX, SB202190, SB203580, CHX+SB202190, CHX+SB203580. At least 3672 cells were counted for each timepoint.

Both p38 inhibitors almost completely prevented cycloheximide from blocking the progression of G2 phase cells into M phase (Figure 6B,C). Whereas only 4 out of 100 cells treated with cycloheximide alone entered M phase, 83 out of 100 SB202190- and 77 out of 92 SB203580-treated cells did enter M phase in the presence of cycloheximide (Figure 6C). For cells treated with cycloheximide alone, the probability of entering M phase was near-zero unless the cells were treated late in G2 phase (Figure 6C,D), as shown earlier (Figure 3E); however, the probability of entering M phase for cells treated with cycloheximide plus either SB202190 or SB203580 was 10-20% for cells treated at the start of G2 phase and rose to near 100% for cells treated 2 h after the start of G2 phase (Figure 6C,D). Overall 50 out 95 cells treated with SB202190 plus cycloheximide and 59 out of 91 cells treated with SB203580 plus cycloheximide exhibited a normal G2 phase duration (within two standard deviations of the DMSO treated control, Figure 6 – figure supplement 1). However, a significant fraction of the cells treated early during G2 phase had prolonged G2 durations, whereas cells treated later in G2 phase mostly exhibited a normal G2 duration (Figure 6–figure supplement 1B). The combination of cycloheximide plus SB202190 or SB203580 caused a small increase in the mitotic index followed by a slow decline to lower levels but rescued the sharp decline in mitotic cells observed in cycloheximide-treated cultures (Figure 6E and Figure 6–figure supplement 2B). The structurally similar but inactive compound SB202474 did not prevent cycloheximide from blocking M-phase entry (Figure 6–figure supplement 2). Similar results were obtained in HeLa and hTERT-RPE1 cells (Figure 6–figure supplement 2C,D).

Thus, the p38 MAPK inhibitors SB202190 and SB203580 allow the majority of the cycloheximide-treated G2-phase cells to progress into M phase. This suggests that p38-mediated checkpoint effects, rather than a lack of protein synthesis per se, are principally responsible for the arrest of cycloheximide-treated G2-phase cells.

Cells treated with SB202190 or SB203580 alone generally progressed through G2 phase normally, although about 20% of cells showed a prolonged G2 phase (Figure 6–figure supplement 1A) and a few cells (4/92 and 2/90) failed to enter M phase (Figure 6C). Thus, in normal, unperturbed cells, p38 has relatively little effect on G2 duration and M phase entry in cells that have not been treated with cycloheximide.

### Protein synthesis during G2 phase is required for normal mitotic progression

So far, we have shown that inhibition of either Wee1/Myt1 or p38 can overcome a cycloheximide-induced arrest and allow G2 phase cells to progress into mitosis in the absence of protein synthesis. However, the requirements to enter mitosis and to successfully progress through mitosis might differ (see, e.g. Gavet and Pines (22)). Consistent with this notion, we had already observed that cells treated with the Wee1/Myt1 inhibitor during S phase could enter mitosis (with little or no G2 phase) but then exhibited pronounced mitotic errors (Figure 5). In addition, a few cells treated with cycloheximide plus PD0166285 during G2 phase exhibited very long mitoses (Figure 4D), as did some cells treated with cycloheximide plus either of the p38 inhibitors (Figure 6C). Accordingly, we examined the duration of mitosis in cells treated in G2 phase with cycloheximide plus either the Wee1/Myt1 inhibitor or one of the p38 inhibitors. A substantial fraction of the cells treated with PD0166285 with or without cycloheximide exhibited a protracted mitosis (47% for PD-treated cells and 58% for cells treated with PD plus cycloheximide, Figure 7A), taken here as mitosis longer than two standard deviations above the average duration of mitosis in DMSO-treated cells. Thus, PD- or PD-plus cycloheximide treated cells that successfully entered mitosis were, nevertheless, delayed in their progression through M phase. These results confirm previous findings that Wee1/Myt1 inhibition extends the mitotic duration (45). The duration of mitosis was a dose-dependent function of the PD0166285 concentration (Figure 7B; cf. Figure 4B). Moreover, the greater the shortening of G2 phase, the greater the delay in M phase (Figure 7–figure supplement 1).

**Figure 7.**
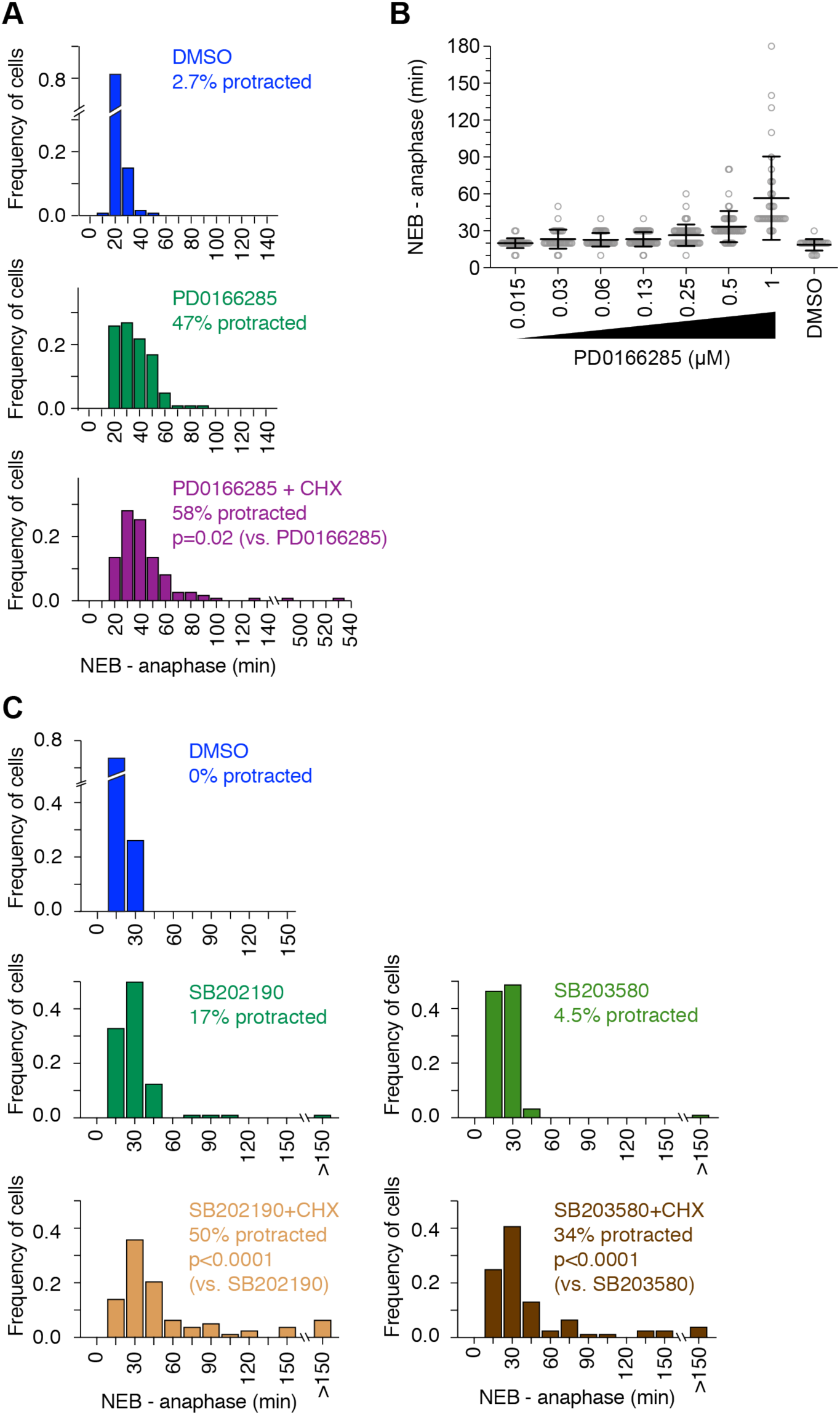
Protein synthesis during G2 phase is required for normal mitotic progression. (**A**) Frequency distribution of mitotic durations (measured from nuclear envelope breakdown (NEB) to anaphase onset) of cells treated with DMSO (n=113),1 *µ*M PD0166285 (n=100) or CHX plus 1 *µ*M PD0166285 (n=110) during G2 phase. Cells were considered to exhibit a protracted mitosis if the mitotic duration was longer than two standard deviations above the average duration of mitosis in DMSO-treated cells. P-values were calculated using a nonparametric Mann-Whitney test. (**B**) Mitotic duration was measured for cells that progressed through G2 phase and entered mitosis in the presence of DMSO or different concentrations of PD0166285. Mitotic duration increased with higher concentrations of PD0166825 (and shorter G2 duration, see Figure 4B). (**C**) Frequency distribution of mitotic durations of cells treated with DMSO (n=99), SB202190 (n=88), SB203580 (n=90), CHX+SB202190 (n=78) or CHX+SB203580 (n=77) during G2 phase. P-values were calculated using a nonparametric Mann-Whitney test.

For the p38 inhibitors SB202190 and SB203580, the drugs had some effect on the duration of mitosis even in the absence of cycloheximide; 17% (SB202190) and 4.5% (SB203580) of the drug-treated cells exhibited a protracted mitosis, versus 0% for the DMSO-treated controls (Figure 7C). This suggests that p38 function may contribute to M-phase progression in at least a subset of cells. A greater proportion of cells treated with either of the inhibitors plus cycloheximide exhibited a protracted mitosis (50% and 34% respectively), suggesting that the protein synthesis that normally occurs during G2 phase helps cells to progress through M phase in a timely fashion.

## Discussion

Here we have used live-cell imaging to confirm the decades-old observation (1, 2) that the protein synthesis inhibitor cycloheximide prevents G2 phase cells from entering mitosis (Figure 3). Based on logistic regression analysis, the point of no return–the time at which the cell becomes refractory to cycloheximide treatment–occurs only at the end of G2 phase (Figure 3). On a cell biological level this timing approximately corresponds to when the antephase checkpoint is silenced (54–56). On a biochemical level, this is about when the activity of cyclin B1-Cdk1 rises to maximal levels (12, 15, 22). It is possible that all three of these phenomena are manifestations of the flipping of the bistable Cdk1/PP2A switch from its interphase to its M-phase state.

The cycloheximide effect appears not to be due to the inhibition of protein synthesis per se, but rather to the activation of p38 MAPKs. Accordingly, the p38 inhibitors SB202190 and SB203580 largely restored mitotic entry in cycloheximide-treated cells (Figure 6 and Figure 6–figure supplement 1 and 2). Likewise, the Wee1/Myt1 inhibitor PD0166285 allowed cycloheximide-treated cells to progress into mitosis, consistent with the hypothesis that the effects of the p38 inhibitors are ultimately mediated by the Cdc25 and/or Wee1/Myt1 proteins (Figure 4). Taken together, these results suggest that G2 phase cyclin synthesis, and G2 phase protein synthesis in general, is not strictly required for timely progression into M phase.

These findings also suggest that the accelerating accumulation of cyclin B1 that normally begins at about the onset of G2 phase is not the trigger for mitosis, or at least not the only trigger, since normal G2 durations can be seen in the absence of such protein synthesis.

This conclusion fits well with loss of function studies that show that cyclin A2 synthesis but not cyclin B1 synthesis is required for mitotic entry (57–59). Likewise, the current findings fit well with the observation that cyclin B1 overexpression has little or no effect on cell cycle dynamics (60).

What then is the trigger for mitosis? One possibility is that the cessation of some low but non-zero levels of ATR- or ATM-mediated checkpoint signaling (11, 57) at the S/G2 boundary might set into motion a signal transduction process that leads to inactivation or degradation of Wee1/Myt1 (61–63) and activation of Cdc25. Consistent with this hypothesis, we observed that the duration of G2 phase is a sensitive function of the basal level of Wee1/Myt1 activity (Figure 4B). Another possibility is that the translocation of cyclin A2 from the nucleus to the cytoplasm, an event that occurs during late S phase, may initiate the events that lead to mitotic entry (64, 65). A third possibility is that the Bora-Aurora A-Plk1 pathway is the critical trigger (18, 66). How exactly cells integrate the different signaling pathways in order to decide whether or not to enter mitosis remains an important, open question in somatic cell cycle regulation.

Cycloheximide-treated cells rescued by p38 MAPK inhibition, and cells entering mitosis precociously due to Wee1/Myt1 inhibition, did take longer to progress through and exit mitosis (Figure 7). In both cases, these cells would be expected to enter mitosis with lower cyclin B1 levels than normal. The lower cyclin B1 levels could result in all or some mitotic substrates being phosphorylated more slowly, resulting in the observed mitotic delays.

G2 phase protein synthesis, or cyclin B1 synthesis more specifically, appears to represent “just-in-time” preparation for the next phase of the cell cycle. This concept, borrowed from supply chain management, has been proposed to apply to protein synthesis and complex assembly in the bacterial cell cycle (67). Even though the proteins involved in the bacterial cell cycle bear little resemblance to those that regulate the eukaryotic cell cycle, perhaps this concept applies to both regulatory systems.

## Materials and Methods

### Cell culture

MCF10A human mammary epithelial cells were a kind gift from Sabrina Spencer and were cultured in growth medium DMEM:F12 (Gibco, #11320-033) containing 5% horse serum (Gibco, catalog number 16050114), 20 ng/ml EGF (PreproTech, AF-100-15), 0.5 mg/mL hydrocortisone (Sigma-Aldrich, H0888-1g), 100 ng/ml choleratoxin (Sigma-Aldrich, C8052-2mg) 10 *µ*g/ml insulin (Sigma-Aldrich, I1882-100mg), 1% penicillin, and 1% streptomycin (both from Life Technologies, catalog number 15140-122) as described previously (41, 42). HeLa cells (CCL-2) were purchased from the ATCC and cultured in Dulbecco’s modified Eagle medium containing high glucose and pyruvate (Invitrogen, #11995-073) supplemented with 10% fetal bovine serum (Axenia Biologix, #F001), 1% penicillin, 1% streptomycin, and 4 mM L-glutamine (all from Gemini Bio-Products, #400-110). HEK 293T cells were obtained from ATCC (CRL-3216) and cultured in the same medium as HeLa. hTERT RPE-1 cells were obtained from ATCC (CRF-4000) and cultured in DMEM:F12 (Gibco, catalog number 11320-033), 10% fetal bovine serum (Axenia Biologix, #F001), 0.01 mg/ml hygromycin B (Invitrogen, 10687-010), 1% penicillin, and 1% streptomycin. All cells were maintained at 37°C and 5% CO2 and discarded after passage 25.

### Stable cell lines

To obtain MCF10A cells stably expressing eYFP-PCNA, we sub-cloned eYFP-PCNA from the eYFP-PCNA construct (43) into the pTRIP-EF1α lentiviral transfer vector (68), kindly provided by Ed Grow at Stanford University, by using the *Xba*I and *Bam*HI restriction sites. To make lentivirus, we incubated 1 ml Opti-MEM and 36 μl FuGENE6 for 5 min at room temperature, then added 10 μg pTRIP-EF1α-eYFP-PCNA, and 6.6 μg pCMVΔR8.74, 3.3 μg pMD.G-VSVG, and 3.3 μg pRev (all kindly provided by Ed Grow), and incubated for 30 min. We used this to transfect HEK 293T cells in Opti-MEM for 6 h at 37°C and 5% CO2 and then exchanged with fresh Opti-MEM. We harvested medium containing virus 48 h, 72 h, and 96 h later, filtered out cell debris with a sterile 0.45-μm filter (Millipore), concentrated by centrifuging for 20 min at 3600 rpm in Amicon-Ultra 15 Filter Units with a 100,000 kDa MW cutoff (Millipore), and froze down at −80°C. To transduce MCF10A cells, we added concentrated virus and 5 μg/ml polybrene (Sigma-Aldrich) to MCF10A cells in growth media, incubated for 24 h, and replaced with growth media. After culturing cells for 5 more days, we sorted for eYFP-positive cells by fluorescence-activated cell sorting. To obtain MCF10A cells stably expressing eYFP-PCNA and histone H2B-mCherry or histone H2B-mTurquoise we made lentivirus as above with histone H2B-mCherry or histone H2B-mTurquoise sub-cloned into the CSII-EF lentiviral transfer vector (69). We used it to transduce MCF10A cells stably expressing eYFP-PCNA and then sorted for cells positive for both fluorescent proteins using fluorescence-activated cell sorting. The hTERT RPE-1 cells stably expressing eYFP-PCNA and histone H2B-mTurquoise cells were produced in the same manner.

To obtain HeLa cells stably expressing eYFP-PCNA, we linearized the eYFP-PCNA construct (43) by incubating with *Fsp*I (New England Biolabs) and purifying with ethanol precipitation. We co-transfected linearized eYFP-PCNA and linearized hygromycin marker (Clontech) with FuGENE6 (Promega) at a ratio of 1 μg eYFP-PCNA to 0.1 μg hygromycin marker to 6 μl FuGENE6 according to the manufacturer’s instructions except that we washed cells with Opti-MEM (Invitrogen), transfected cells in Opti-MEM, and incubated in Opti-MEM for 5 h at 37°C and 5% CO_2_ before replacing with growth medium. We split cells 48 h later and after 25 more h added 400 μg/ml hygromycin B (Invitrogen). We picked colonies 11 days later using cloning rings and expanded a clone that had correct PCNA localization in the nucleus and could form PCNA foci.

### Chemical inhibitors

PD0166285 was generously provided by Pfizer and later purchased from EMD Millipore (#513028) and stored frozen as a 25 mM stock in DMSO (Sigma-Aldrich) and used at a final concentration of 1 *µ*M if not specified differently. Cycloheximide was purchased from Sigma-Aldrich, stored frozen as a 10 mg/ml stock in water and used at a final concentration of 10 *µ*g/ml. SB202474 (EMD Millipore, #559387) and SB202190 (Sigma, S7067) were stored frozen as 50 mM or 10 mM stock solutions in DMSO. SB203580 (EMD Millipore, #559387) was stored frozen as a 50 mM stock solution. SB202474, SB202190 and SB203580 were used at a final concentration of 50 *µ*M if not specified differently.

### Live-cell time-lapse microscopy and image analysis

Cells were seeded into 96-well plates (Costar) or collagen-coated (PureCol, Advanced BioMatrix, #5005-1ML) 96-well glass bottom plate (Cellvis, P96-1.5H-N) the day before microscopy at such a density that they were sub-confluent even at the end of the experiment. To prevent drying, each well of the plate contained between 100 and 200 μl of growth medium. Images were taken in 10 min or 15 min intervals, depending on the needs of the experiment, on the ImageXpress Micro System Standard Model (Molecular Devices) controlled by the MetaXpress 5.1 software (Molecular Devices) using the 10X objective (NA = 0.3, Plan Fluor) or the 20X objective (NA = 0.45, Plan Fluor ELWD). Cells were kept alive inside the microscope in a humidified chamber at 37°C and 5% CO_2_. We used the YFP-LIVE filter cube for imaging eYFP-PCNA, the CFP-LIVE filter cube for imaging histone H2B-mTurquoise, the HcRED-LIVE filter cube for imaging histone H2B-mCherry. Combined exposure through all the filter cubes did not exceed 700 msec per frame. We used 4x gain and 1×1 or 2×2 binning.

Raw TIFF images were exported using the MetaXpress 5.1 software (Molecular Devices) and collated into time series by well and site using a script written in MATLAB (MathWorks) or Python. Cells were tracked manually and each relevant change in a fluorescent reporter (PCNA focus disappearance, etc.) was recorded in Excel (Microsoft). For some analyses we used a graphical user interface written in LabView (National Instruments) that recorded the frame number and cell coordinates by responding to a mouse click and exported results to Excel (Microsoft). To allow features like PCNA foci to be easily perceived, images were typically min-max adjusted, and we sometimes allowed the H2B-mCherry image to be saturated in the mitotic stages in order to allow the low intensity H2B-mCherry signal in interphase to be perceivable. A custom-written Matlab script provided by Tobias Meyer’s laboratory was used to count the total number of cells in every time frame in order to calculate mitotic indices.

### Immunoblotting and antibodies

2 ml of MCF10A cell suspension at a cell concentration of 1.5 x 10^5^ cells/ml was seeded into a 6-well plate (Falcon, #353046) and grown for 48 h at 37°C. The medium was exchanged and the cells were grown for another 6 h. An equal volume of medium containing the prediluted inhibitors was then added to the cells, and the cells were incubated for 30 min (for PD0166285) or 6 h (for cycloheximide and cycloheximide plus either SB202190 or SB203580). The medium was then removed, cells were washed twice with cold PBS and lysed with lysis buffer (20 mM Tris-HCl pH 7.4, 150 mM NaCl, 1% NP-40, 1 mM EDTA, 1x Phosstop #4906845001, 1x cOmplete #11873580001). Samples were boiled in SDS PAGE sample buffer and analyzed by SDS PAGE and immunoblotting. The following antibodies were used: rabbit α-Cdk1 phospho-Tyr15 (Cell Signaling Technology, #9111L), mouse α-Cdk1 (Santa Cruz Biotechnology #SC-54), mouse α-tubulin (Santa Cruz Biotechnology #SC-32293), rabbit α-HSP27 phospho-Ser82 (Cell Signaling Technology, #2401), mouse α-HSP27 (Cell Signaling Technology, #2402), rabbit α-p38 MAPK (Cell Signaling Technology, #9212) and rabbit α-p38 MAPK phospho-Thr180/Tyr182 (Cell Signaling Technology, #9211).

### Statistical analysis

Logistic regression analysis is a method for estimating how the probability of a binary outcome—in our case, whether a cell does or does not ultimately progress into mitosis— varies as a function of time and treatment conditions. The underlying assumption is that the odds of progressing into mitosis scale multiplicatively with time, which means that the time course data should be approximated by a logistic function, with parameters for the steepness and time of the transition from low to high probability. We binarized cell outcomes (i.e., the cell either did or did not progress into mitosis within the average G2 phase duration plus two standard deviations of the DMSO-treated control), plotted the fraction of cells that attained the outcome as a function of the time of drug addition, and fitted the data to a logistic function using the LogitModelFit command in Mathematica 10. The 95% confidence bands were calculated using code deposited in the Mathematica Stack Exchange (https://mathematica.stackexchange.com/questions/26616/how-can-i-compute-and-plot-the-95-confidence-bands-for-a-fitted-logistic-regres).

Further statistical analyses were performed using Graphpad Prism 8.0.2.

All experiments except the immunoblot analyses have been performed in at least two biological replicates – meaning that cells were freshly plated, imaged and independently treated with the respective drugs – of which usually one representative experiment is shown (sometimes we show both).

## Acknowledgments

We thank Mary Teruel, Michael Zhao and the Stanford High Throughput Biosciences Facility for help with the ImageExpress microscopy; We thank Arne Lindqvist, Tim Stearns, Tobias Meyer, Karlene A. Cimprich, Aaron F. Straight, Frederick G. Westhorpe, Whitney L. Johnson, and Bradley T. French for helpful discussions. We are indebted to Sabrina L. Spencer, Ed Grow, Tony Yu-Chen Tsai, Paul G. Rack, M. Cristina Cardoso, and Michael W. Davidson for sharing reagents. We acknowledge help with LabView from Taru Roy, help with image analysis from Colin J. Fuller, and help with MATLAB from Feng-Chiao Tsai, Sabrina Spencer, Jeremy B. Chang, Graham A. Anderson, and Tony Yu-Chen Tsai. We are grateful to Pfizer for providing PD0166285. We thank the Stanford FACS Facility and the Stanford High-Throughput Biosciences Facility for technical assistance; and Silke Hauf, Oshri Afanzar, Xianrui Cheng, Yuping Chen, William Huang, Shixuan Liu, and Connie Phong, for comments on the manuscript. The work was supported by grants from the National Institutes of Health (R01 GM046383 and R35 GM131792, J.E.F.), a Stanford Graduate Fellowship (S.L.), the Stanford Training Grant in Chemical and Molecular Pharmacology (T32 GM067586, S.L.), and a postdoctoral fellowship from the German Research Foundation (KA 4476/1-1, J.K.).

---

**Figure 4–figure supplement 1.**
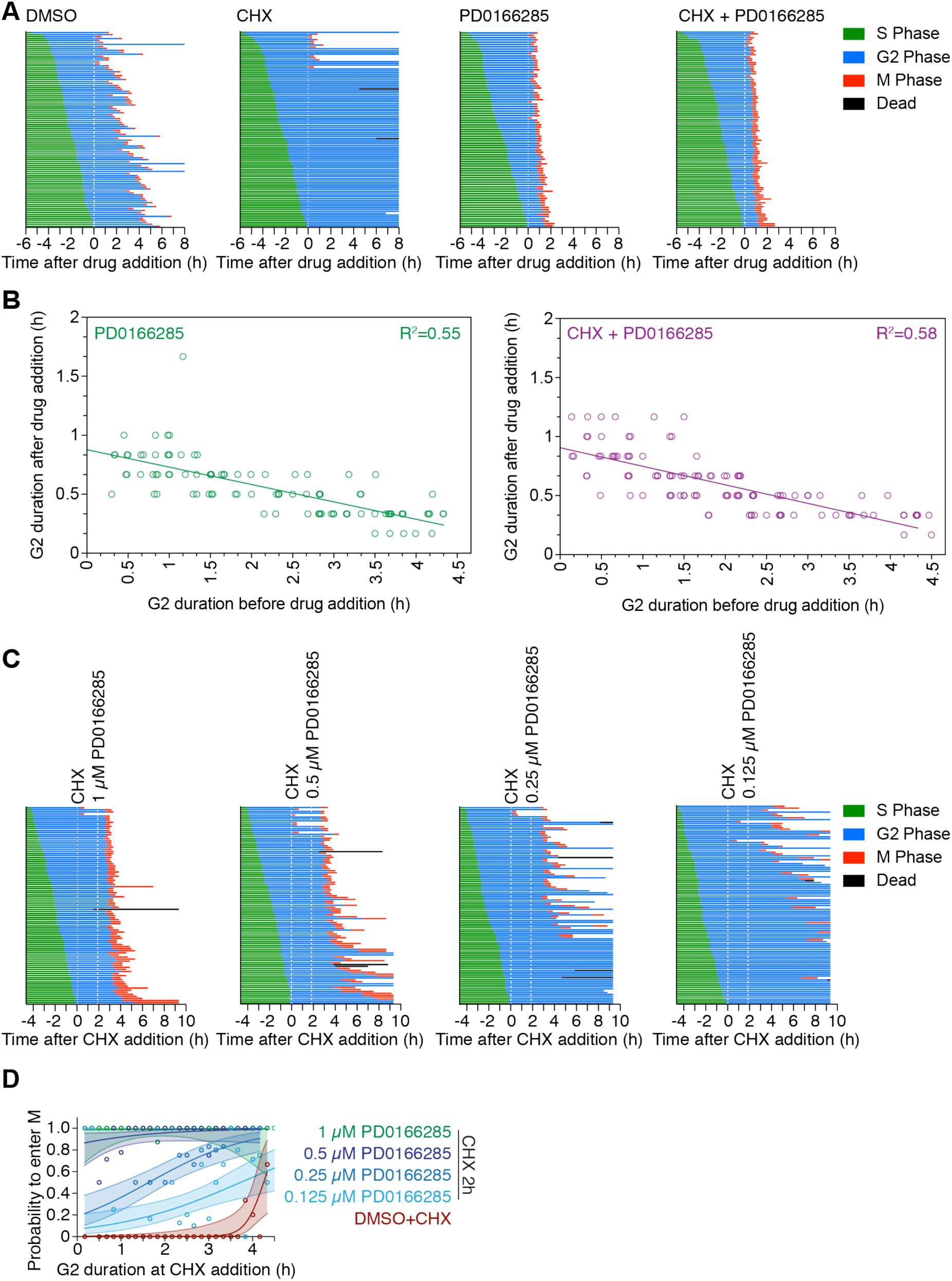
Wee1/Myt1 counteract pro-mitotic activities accumulating during G2 phase. (**A**) Cell fate trajectories of MCF10A cells expressing H2B-mCherry and eYFP-PCNA treated with DMSO (n=100), cycloheximide (n=107), PD0166285 (n=85) or cycloheximide plus PD0166285 (n=103). Biological replicate of data shown in Figure 4D. (**B**) The time between drug addition (1 *µ*M PD0166285 or 1 *µ*M PD0166285 plus cycloheximide) and mitotic entry as a function of how much time cells had spent in G2 at the time of drug addition. Cells early in G2 phase required more time to enter mitosis than cell late in G2 phase after Wee1/Myt1 inhibition. Circles show individual cells. In order to better differentiate data points in the x-dimension minimal random noise was added to each data point. Lines correspond to linear least square regression fits. (**C**) Cell fate trajectories of MCF10A cells expressing H2B-mCherry and eYFP-PCNA treated with cycloheximide for 2 h before the addition of different concentrations of Wee1 inhibitor PD0166285 (n= 95 for 1 *µ*M PD0166285, n=103 for 0.5 *µ*M PD0166285, n=107 for 0.25 *µ*M PD0166285, n=104 for 0.125 *µ*M PD0166285). (**D**) Probability of a cell to enter mitosis as a function of how long the cell has already been in G2 phase at the time of cycloheximide addition for the experiment shown in (C). Circles indicate the frequency of mitotic entry for any given G2 duration, solid lines show the logistic fit for the data and lightly colored areas indicate the covariance of the fit.

**Figure 6–figure supplement 1.**
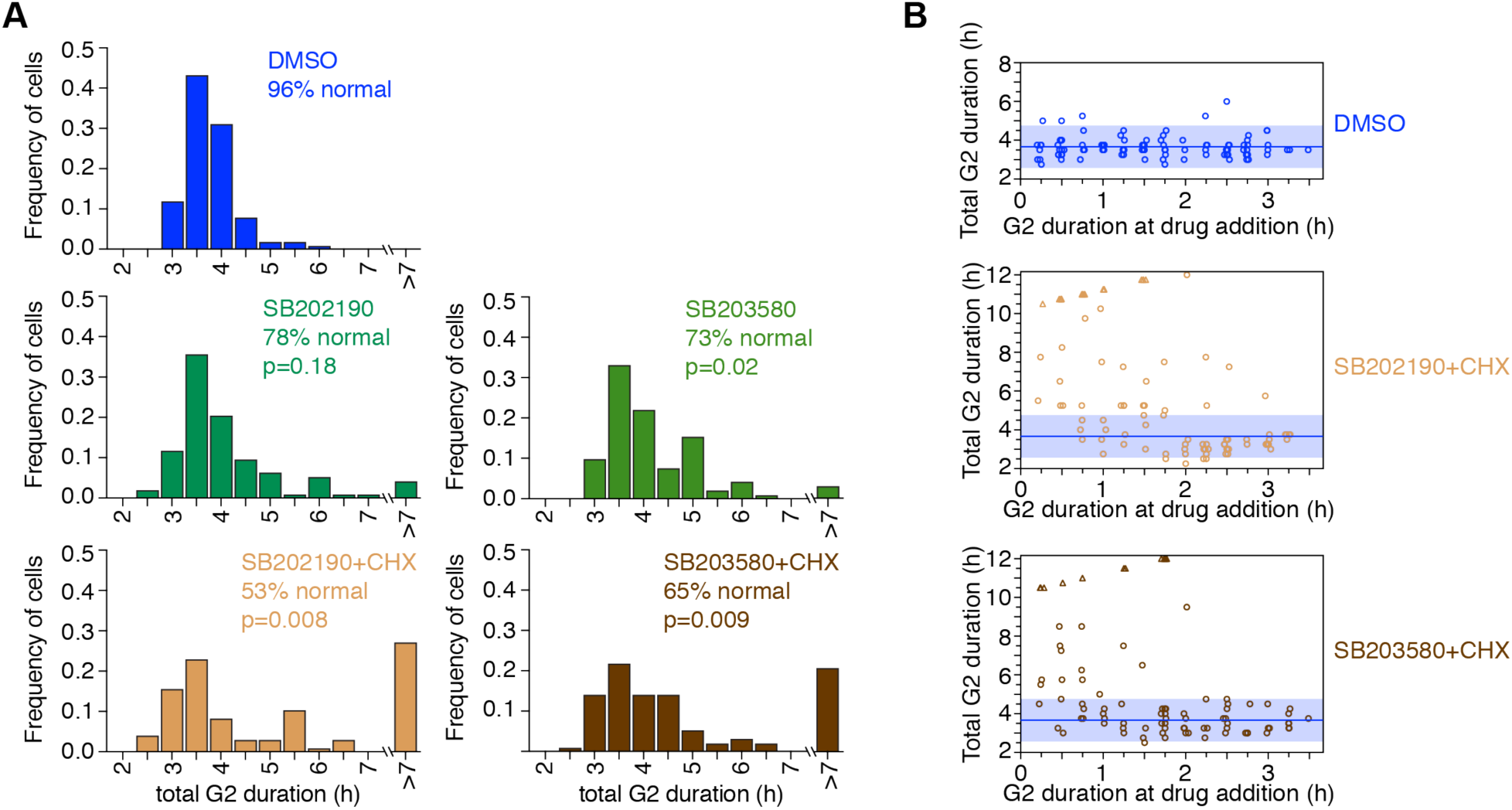
The majority of cells exhibit a normal G2 duration when treated with cycloheximide plus p38 inhibitors. (**A**) Frequency distribution of the total duration of G2 phase of cells treated with DMSO (99), SB202190 (n=92), SB203580 (n=95), cycloheximide plus SB202190 (n=90) or cycloheximide plus SB203580 (n=91). P-values were calculated using a nonparametric Mann-Whitney test. (**B**) Total G2 duration plotted against the time a cell had spent in G2 at the time of drug addition for cells treated with DMSO, or cycloheximide plus either SB202190 or 203580. Blue, solid line depicts the mean G2 duration for the DMSO control and the light blue, shaded area marks the area that falls within two standard deviations of the mean. Circles depict cells that entered mitosis during the experiment, triangles depict cells that were still in G2 at the end of the experiment. In order to better differentiate data points in the x-dimension minimal random noise was added to each data point.

**Figure 6–figure supplement 2.**
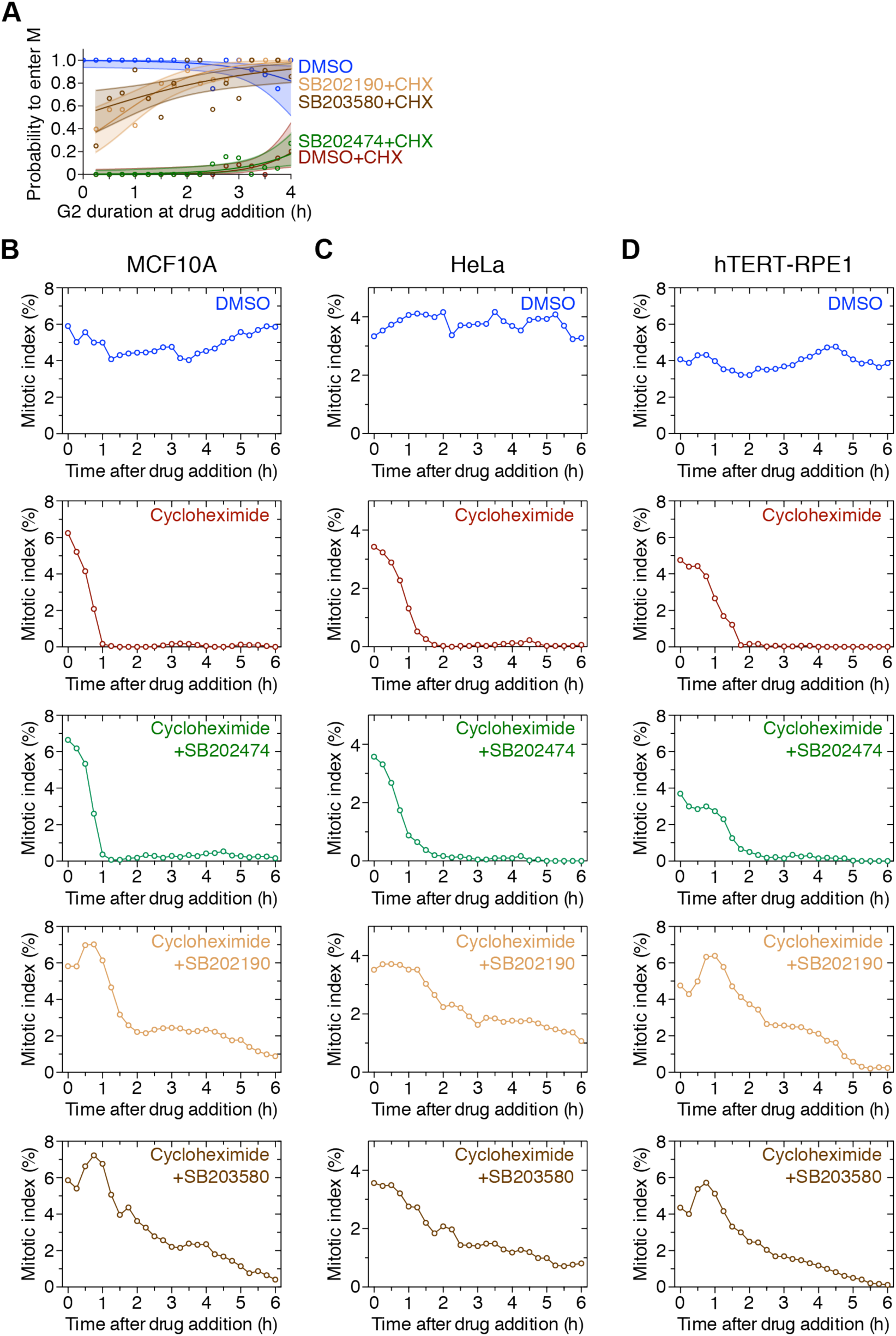
p38 inhibition allows cells to enter mitosis in the presence of cycloheximide. (**A**) Probability of a cell to enter mitosis as a function of how long the cell has already been in G2 phase at the time of drug addition for MCF10A cells expressing H2B-mCherry and eYFP-PCNA treated with DMSO, DMSO plus cycloheximide, cycloheximide plus either p38-inhibitor SB202190 or SB203580, or cycloheximide plus the structural analog SB202474. (**B-D**) Mitotic indices for MCF10A cells (B), HeLa (C) and hTERT-RPE1 (D) cells expressing H2B-Turquoise and eYFP-PCNA cells treated with DMSO, DMSO plus cycloheximide, cycloheximide plus either p38-inhibitor SB202190 or SB203580, or cycloheximide plus the structural analog SB202474. For each timepoint at least 3605 (MCF10A), 3018 (HeLa), and 2506 (hTERT-RPE1) cells were counted.

**Figure 7–figure supplement 1.**
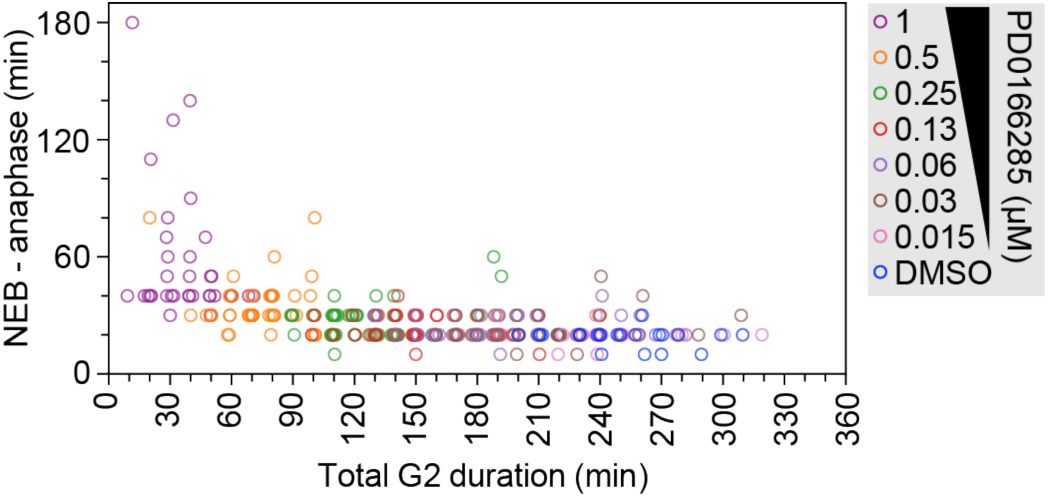
Inverse relationship between the duration of G2 phase and the duration of M phase in PD0166285-treated cells. Mitotic timing (measured as the time from nuclear envelope breakdown to anaphase onset) for cells that were treated with cycloheximide for 2 h before the addition of 1 *µ*M PD0166285 as a function of how much time the cell had spent in G2 phase at the time of cycloheximide addition. In order to better differentiate data points in the x-dimension minimal random noise was added to each data point. Note that cells that had spent less time in G2 phase at cycloheximide addition, required more time to progress into anaphase.

